# Seasonality controls the predictive skills of diatom based salinity transfer functions

**DOI:** 10.1101/341446

**Authors:** Alejandra Goldenberg Vilar, Timme Donders, Aleksandra Cvetkoska, Friederike Wagner-Cremer

## Abstract

The value of diatoms as bioindicators of contemporary and palaeolimnological studies through transfer function development has increased in the last decades. While they represent a tremendous advance in (palaeo) ecology, these models also leave behind important sources of uncertainties that are often ignored. In the present study we tackle two of the most important sources of uncertainty in the development of diatom salinity inference models: the effect of secondary variables associated to seasonality and the comparison of conventional cross-validation methods with a validation based on independent datasets. Samples (diatoms and environmental variables) were taken in spring, summer and autumn in the freshwater and brackish ditches of the province of North Holland in 1993 and sampled again different locations of the same province in 2008-2010 to validate the models. We found that the abundance of the dominant species significantly changed between the seasons, leading to inconsistent estimates of species optima and tolerances. A model covering intra-annual variability (all seasons combined) provides averages of species optima and tolerances, reduces the effect of secondary variables due to the seasonality effects, thus providing the strongest relationship between salinity and diatom species. In addition, the ‘all-season’ model also reduces the edge effects usually found in all unimodal-based calibration methods. While based on cross-validation all four models seem to perform relatively well, a validation with an independent dataset emphasizes the importance of using models covering intra-annual variability to perform realistic reconstructions.

## Introduction

Diatom assemblage changes provide an excellent basis for inferring environmental changes from seasonal scales to decadal or centennial scales given their sensitivity to a broad variety of habitat parameters. Available transfer functions cover nutrient status of freshwater bodies, temperature, as well as salinity dynamics [1-5]. Diatom-based models to infer salinity or tidal height have recently become an increasing focus in transfer function development as a potential tool in sea level reconstruction efforts [6-8]. These models require extremely high predictive precision given the often subtle changes they need to quantify. However, the application of transfer functions to model a single variable (e.g. salinity) is problematic under multiple response triggers [9, 10]. The effect of secondary gradients on model performance and importantly on the forthcoming reconstructions, has yet received surprisingly little attention. Quantitative reconstructions are a form of space-for-time substitution [11], and as such require that the co-variation between the modelled variable and potentially confounding underlying ecological factors is constant in both time and space. In most situations this is difficult to test, but nonetheless a highly unrealistic assumption. Firstly, this inherent problem of quantitative reconstructions is especially evident in relation to seasonality (referring to biological and chemical changes occurring in continental waters according to the different seasons in temperate climates). In diatom populations, seasonal succession is characterized by changes in nutrient concentrations, light [12], thermal stratification and predator–prey relationships [13, 14]. In addition, diatom communities follow distinct seasonal succession patterns caused by changes in life-history traits [15, 16] and nutrient stoichiometry [17]. Despite the high likelihood that a modern training set selected for building a transfer function will be influenced by the seasonality effects, the strength of such dependences is not yet known, yet. Only a limited number of diatom-based studies have used contemporary data to facilitate interpretation of sediment core in terms intra-annual diatom distribution [18]. The few studies available that have analyzed the effect of seasonality on diatom-based transfer functions generally focused on nutrient variables. However, the effect of many other environmental variables can be obscured by seasonality, including salinity. The potential effect of seasonality is thereby extremely important, given the intra-annual variability of salinity that needs to be separated from the long-term changes. A second limitation in the development of transfer functions is the lack of independent data sets to test the reliability of the reconstructions [19]. Usually the predictive ability of the transfer functions is only assessed by cross-validation methods. If the observations in the calibration set are not independent, because of autocorrelation or other types of pseudo-replication, performance statistics based on cross-validation will be over-optimistic [20]. Therefore, the ideal way of finding unbiased transfer-function performances is the use of an independent test set [20].

In the present study, we tackle two important uncertainty sources in diatom based transfer function development following the recommendations set outlined in Juggins [21]. We evaluate (1) the effect of seasonality when modeling a single variable (salinity) regarding the “true” ecological meaning of statistically significant models and (2) compare model performance between conventional cross validation methods and independent validation dataset. Both approaches aim to detect confounding environmental factors on the primary variable of interest.

## Materials & Methods

### Study area and datasets

The study was conducted in the province of North Holland, the Netherlands. The samples were collected in modified wetlands that have been reclaimed for agriculture representing a system of shallow brackish and freshwater ditches draining water from low-lying areas. Water tables in the three areas are kept within strict limits and the banks are bordered by reed belts dominated by *Phragmites australis*. Most of the diatom species and macrophytes found in these modified wetlands are also found in mesotrophic and eutrophic European lakes. Detailed explanation of the study area is provided in [22].

The dataset for the development of the diatom inference models is referred as ‘training set’ and consists of a total of 96 samples of 32 locations sampled in spring (March), summer (June) and autumn (September) 1993. The samples were collected randomly in the province of North Holland spanning a salinity gradient from 200-9000 mg/l chloride. As a result, salinity represent the most important gradient in the dataset. Water samples were collected every month at the same locations as the diatom samples and the following variables were determined: surface water oxygen, pH, conductivity, chloride, sulphates, transparency (Secchi depth), total nitrogen, total phosphorus and chlorophyll-*a.* A summary of the environmental variables measured is presented in Table 1. In this dataset, 400 diatom valves were counted per sample.

**Table 1.** Descriptive statistics of the environmental variables measured in the training and validation datasets. Temp= temperature; O_2_=dissolved oxygen; tN=total nitrogen; tP=total phosphorus; Cl=chloride; Secchi= Secchi depth; Cond=conductivity; Chl *a*= chlorophyll *a.*

| TRAINING SET |  | Temp. | O<sub>2</sub> | NO<sub>3</sub> | NH<sub>4</sub> | tN | tP | Cl | pH | Secchi | Cond | SO<sub>4</sub> | Chl *a* |
| --- | --- | --- | --- | --- | --- | --- | --- | --- | --- | --- | --- | --- | --- |
|  |  | °C | mg/l | mg/l | mg/l | mg/l | mg/l | mg/l |  | cm | uS/cm | mg/l | ug/l |
| SPRING<br>n=32 | Max | 12.0 | 24.0 | 2.5 | 2.0 | 9.5 | 2.4 | 7700.0 | 9.1 | 100.0 | 32000.0 | 1800.0 | 234.50 |
|  | Min | 5.5 | 6.2 | 0.0 | 0.1 | 2.0 | 0.1 | 230.0 | 7.4 | 10.0 | 9125.0 | 70.5 | 3.00 |
|  | Mean | 9.1 | 11.9 | 0.6 | 0.4 | 5.2 | 0.6 | 2213.9 | 8.4 | 33.4 | 25138.9 | 381.2 | 78.11 |
|  | Median | 9.0 | 11.1 | 0.1 | 0.2 | 4.6 | 0.4 | 1200.0 | 8.5 | 25.0 | 27500.0 | 210.0 | 64.50 |
| SUMMER<br>n=32 | Max | 20.0 | 14.0 | 1.1 | 3.0 | 13.1 | 6.8 | 8700.0 | 9.2 | 90.0 | 32000.0 | 1833.3 | 175.00 |
|  | Min | 11.0 | 1.4 | 0.0 | 0.1 | 1.1 | 0.1 | 190.0 | 7.3 | 5.0 | 9543.3 | 32.0 | 3.00 |
|  | Mean | 17.1 | 8.0 | 0.1 | 0.5 | 4.2 | 1.2 | 2312.2 | 8.3 | 30.0 | 24348.0 | 401.7 | 38.67 |
|  | Median | 17.3 | 8.3 | 0.0 | 0.2 | 3.4 | 0.8 | 1300.0 | 8.3 | 25.0 | 29500.0 | 170.0 | 27.67 |
| AUTUMN<br>n=32 | Max | 18.0 | 15.0 | 2.7 | 5.4 | 10.0 | 2.8 | 7900.0 | 9.5 | 110.0 | 32000.0 | 1700.0 | 167.50 |
|  | Min | 10.0 | 0.8 | 0.0 | 0.1 | 0.6 | 0.1 | 140.0 | 7.3 | 10.0 | 10780.0 | 63.0 | 3.00 |
|  | Mean | 14.5 | 7.3 | 0.3 | 0.7 | 4.2 | 0.9 | 1964.7 | 8.0 | 29.7 | 23868.6 | 377.7 | 40.64 |
|  | Median | 15.5 | 7.5 | 0.1 | 0.3 | 4.1 | 0.8 | 670.0 | 7.9 | 22.5 | 26500.0 | 147.5 | 29.50 |
| VALIDATION SET |  |  |  |  |  |  |  |  |  |  |  |  |  |
| n=90 | Max | 23.5 | 18.2 | 2.0 | 8.1 | 11.6 | 6.6 | 13000.0 | 10.0 | 120.0 | 31360.0 | 1600.0 |  |
|  | Min | 6.4 | 0.0 | 0.1 | 0.0 | 0.9 | 0.0 | 110.0 | 7.4 | 10.0 | 690.0 | 7.0 |  |
|  | Mean | 13.4 | 9.1 | 0.3 | 0.5 | 4.1 | 0.9 | 1316.7 | 8.2 | 37.9 | 4233.4 | 217.9 |  |
|  | Median | 13.8 | 9.0 | 0.1 | 0.1 | 3.5 | 0.7 | 470.0 | 8.2 | 30.0 | 2030.0 | 120.0 |  |

The dataset used for testing the performance of the models (henceforth referred to as “validation dataset”) consists of 90 samples collected in drainage ditches within the framework of a long-term monitoring program in North-Holland carried out by the local water authority during spring 2008-2010 (200-300 valves were identified per sample, which is the standard for routine monitoring in The Netherlands). Fig 1 shows the sampling locations for both datasets. In the validation dataset, water samples for the determination of chloride, nutrients and other environmental variables were collected every month following standardized national protocols accredited by the Dutch Standards Institute [23]. We used chloride average values of the spring months (i.e. March through May) in both training and validation sets, assuming that these values reflect local conditions better than single measurements. A summary of the environmental variables measured in the validation data set can be found also in Table 1.

In both datasets, diatoms were sampled from reed stems (*Phragmites australis*). This emergent macrophyte was the most abundant in the drainage ditches sampled and is the recommended substratum for sampling periphyton in the Netherlands [24]. In this way, differences caused by substratum heterogeneity are avoided. Attached diatoms were prepared using H2O2 digestion and mounted on microscope slides with Permount Mounting Medium (Fischer Scientific, Pittsburgh). Taxonomic identification was based on the volumes of Krammer and Hoffmann [25-29] following standard protocols [30].

**Fig 1.**
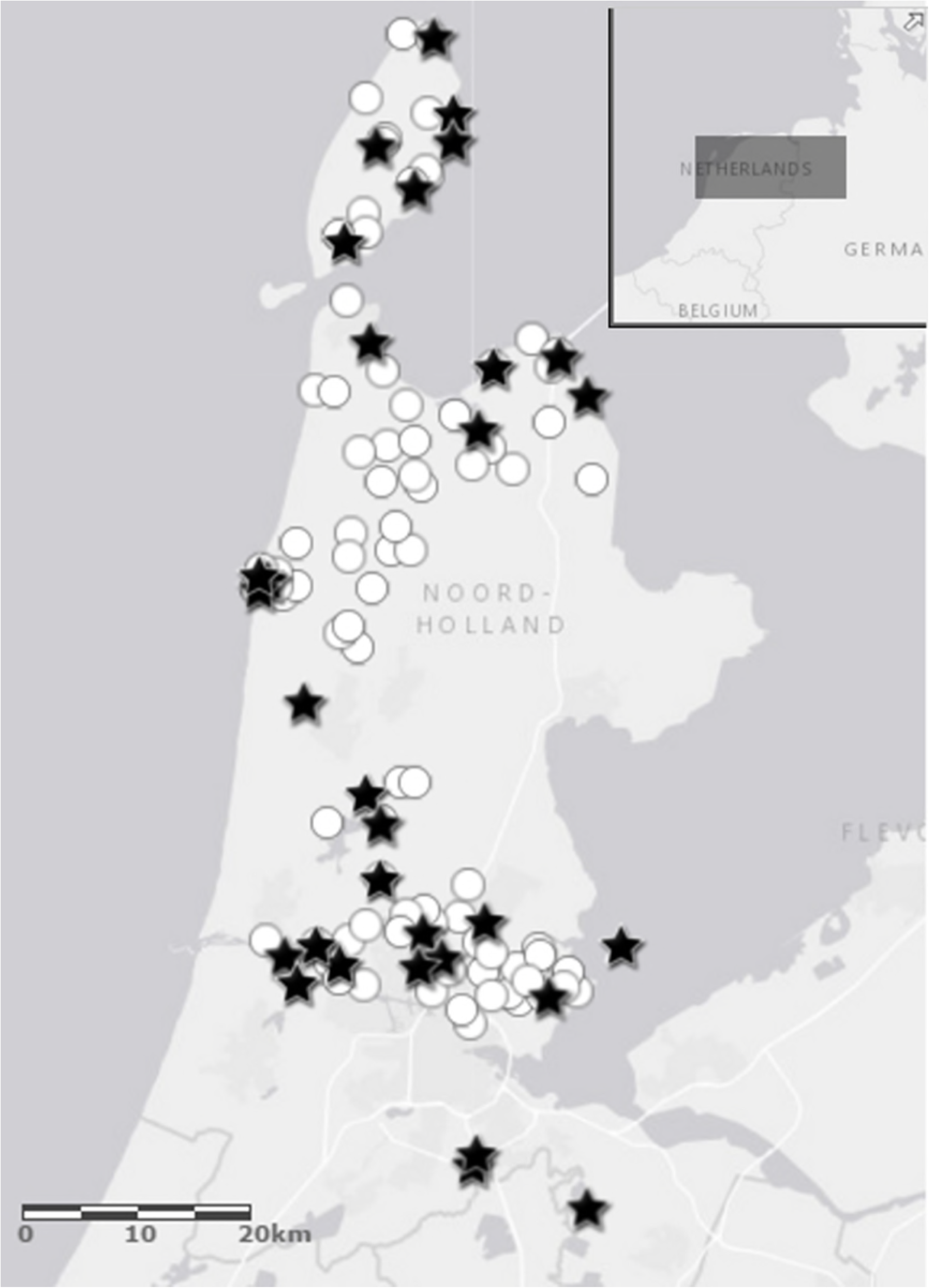
Locations of the training and validation dataset. Training dataset taken in 1993 in spring, summer and autumn (stars) and locations of the validation dataset taken in 2008-2010 in spring (circles) in the province of North Holland (The Netherlands).

## Data analysis

### The effect of seasonality on diatom community structure

We performed two different tests to quantify the effect of seasonality on diatom community structure: 1) permutational multivariate analysis of variance (PERMANOVA) [31] to reflect changes in community composition (species identity and abundances) and 2) analysis of multivariate dispersions [32] that reflects changes in community heterogeneity or beta diversity.

PERMANOVA analyses the variance of multivariate data explained by a set of explanatory factors on the basis of any dissimilarity measure of choice, thereby allowing for a wide range of empirical data distributions. The null hypothesis tested by PERMANOVA is that, under the assumption of exchangeability of the sample units among the groups: “the centroids of the groups are equivalent for all groups” [33]. The test of multivariate dispersions explicitly tests only “the average within-group dispersion (measured by the average distance to group centroid), is equivalent among the groups”. This test is equivalent to the popular Levene’s test in univariate ANOVA but applied to the study of species assemblages by using dissimilarity indices [34].

The statistical significance of multivariate variance components were each tested using 9999 permutations of residuals under a reduced model [31], with an a priori chosen significance level of α=0.05. All multivariate analyses were done on the basis of a Bray–Curtis dissimilarity matrix calculated from square-root transformed percentage abundance data. To visualize multivariate patterns in assemblages across the three seasons, non-metric multi-dimensional scaling (NMDS) was used as an ordination method.

To determine which individual species contributed most to the differences between the seasons we used the species contribution to similarity method (SIMPER), which measures the percentage contribution of each species to average dissimilarity between two groups [35, 36]. One Way Anova was performed to test if environmental variables differed between spring, summer and autumn. These analyses were performed in R statistical package [37].

### The development of diatom based salinity transfer functions

The weighted optima of each species along the salinity gradient (chloride) were determined by averaging all values for each variable from the sites where the taxon occurred, weighted by its abundance at each site. The taxon’s tolerance along each gradient was then calculated as an abundance-weighted standard deviation of the environmental variable [38].

We measured the explanatory strength of salinity as a predictor of diatom assemblage composition by calculating the ratio of the eigenvalue of the first (constrained) CCA axis (λ1) with salinity as a single explanatory variable with the first unconstrained axis (λ2). A value of λ1/λ2 greater than 1 indicates that the variable of interest represents an important ecological gradient in the training set and meets [39] the criterion for a “useful calibration”. To assess the potential confounding effect of other explanatory variables, we performed hierarchical partitioning ordination with the full suite of environmental variables [21].

The WA-PLS regression [40] with “leave one out” cross-validation was used to develop statistical prediction models. This method combines the features of weighted averaging (WA) and partial least squares (PLS) and uses the residual correlation structure in the data to improve the fit between the biological data and environmental data in the training set [41]. The predictive abilities of transfer functions were assessed by examining the relationship between the observed and diatom-inferred values, as well as the observed and jackknife-estimated values of the variables of interest in the training set (r^2^_apparent_ and r^2^_jackknife),_ and evaluation of root mean square error of prediction (RMSEP) [42].

We further evaluated the ability of the model to predict salinity through an independent validation: We used the data from the training set to develop transfer functions and tested their accuracy using the validation dataset. These analyses were performed using C2 version 1.7 software. [43].

## Results

A total of 408 species were identified in the training set of which 179 species covered 98% of all observations and therefore only these were used to develop the models (96 locations in total and 400 individuals counted per sample). In the validation dataset 253 species were identified of which 145 already covered 98% of the data set and were used for the reconstructions (90 locations 200-300 individuals identified per sample). Species percentage data were square-root transformed to reduce the weight of dominant species.

The analysis of PERMANOVA revealed that diatom community composition is affected by seasonality, in terms of overall differences in species abundances: spring-summer *p*=0.001; spring-autumn *p* < 0.001; summer-autumn *p*=0.01 (Table 2 and Fig 2). The species that contributed the most to the differences between the seasons are shown in Table 3. In contrast, the analysis of multivariate dispersions showed that seasonality does not affect beta diversity or community heterogeneity (Table 2). Therefore the two analyses of community structure showed that the seasonal differences in diatom assemblages are related to species abundances and not to species identities. The optima and tolerances of species that contribute the most to community dissimilarity are shown on Fig 3. Either spring or summer models tend to produce the highest optima values, while autumn model lead to the lowest optima. The ‘all-season’ model produces average optima values in comparison to the individual season models.

**Fig 2.**
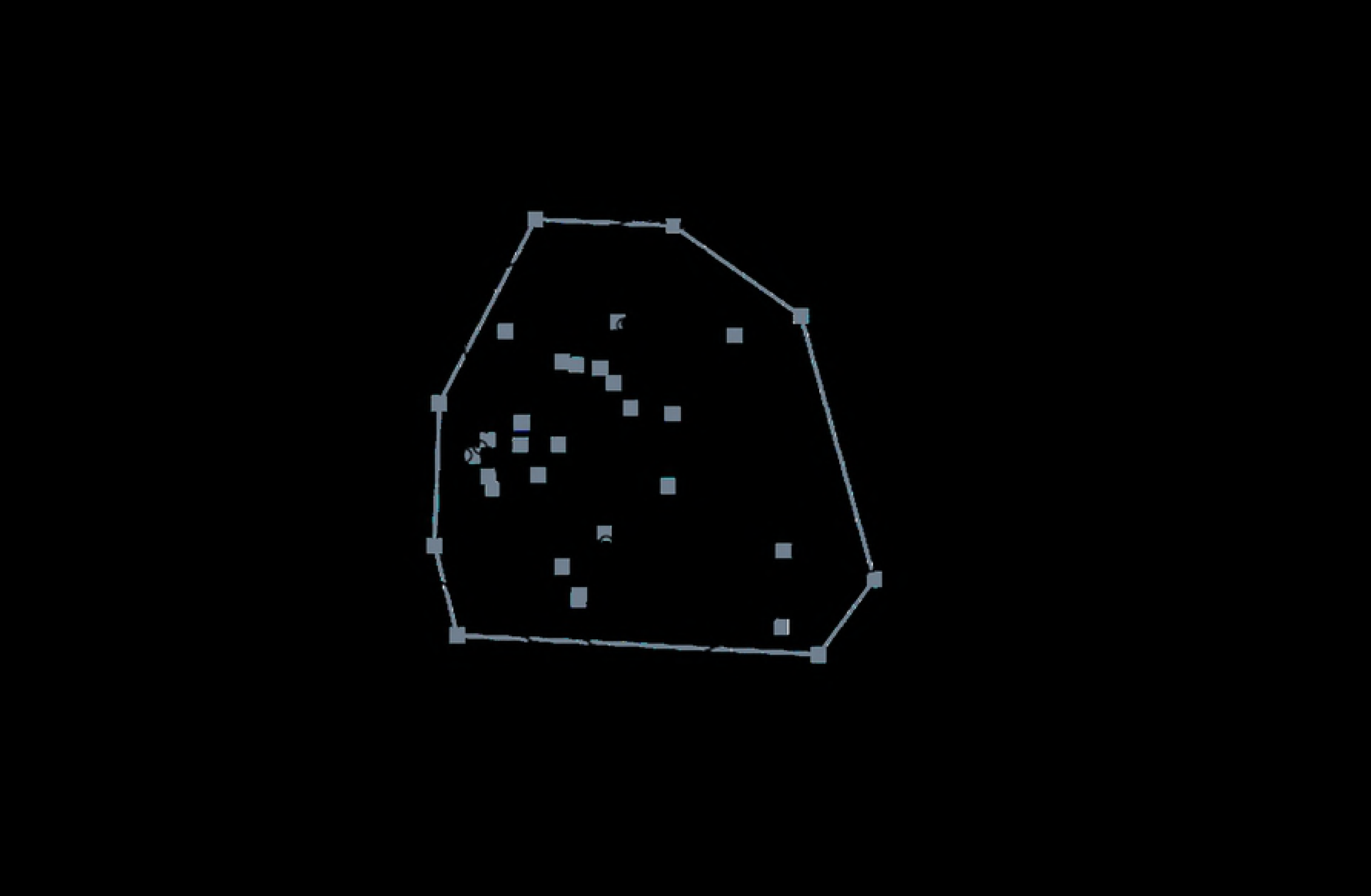
Nonmetric multidimensional scaling ordination diagram of sites based on Bray–Curtis similarity in diatom composition in spring, summer and autumn. Average within group dissimilarity spring: 0.77 (grey squares); summer: 0.83 (open circles); autumn: 0.81 (black circles). The results of the statistical tests of community structure are visible in the graph: the groups differed in their relative position (centroids – PERMANOVA test) and not in the dispersion of the sampling points. (Stress=0.25).

**Fig 3.**
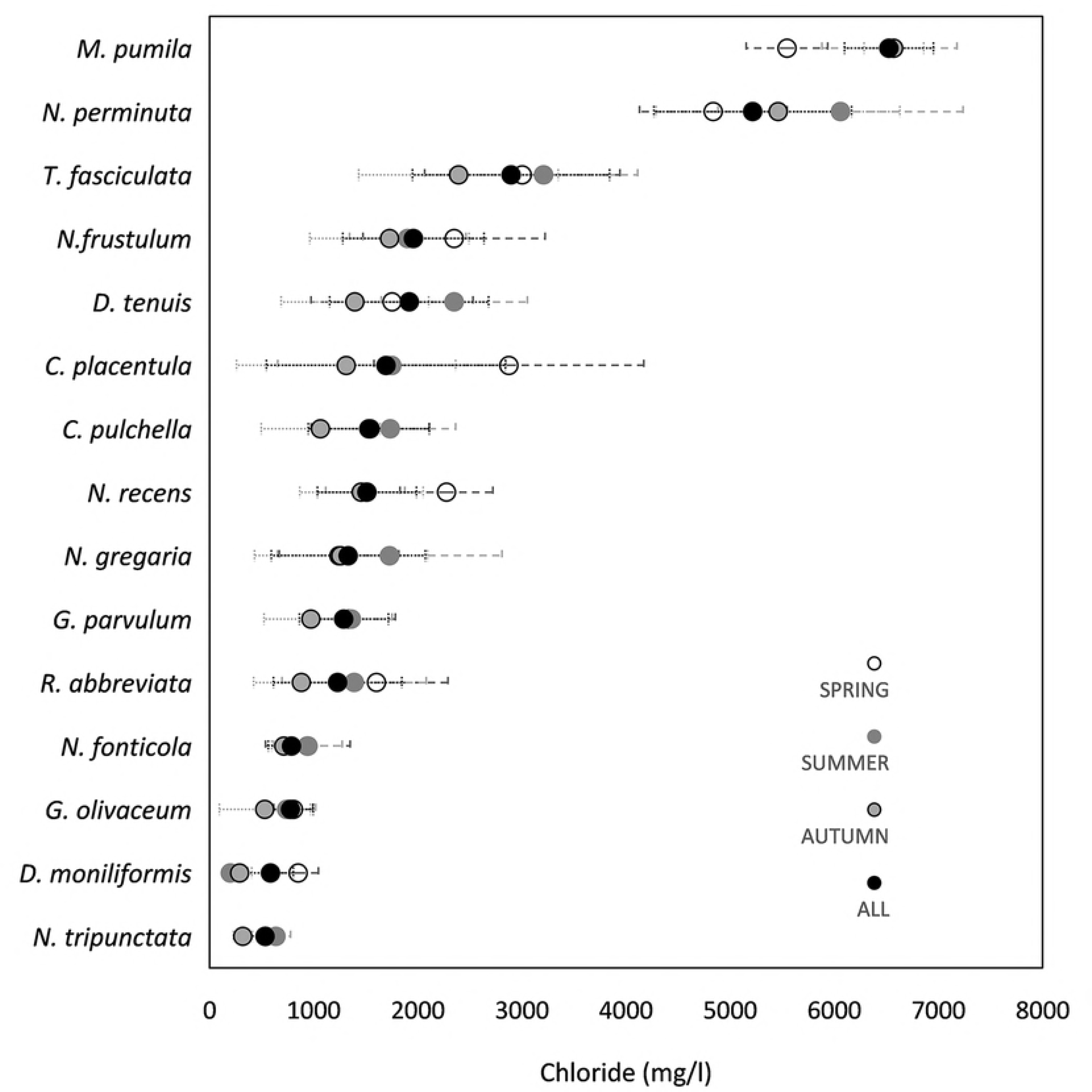
Optima (dots) and tolerances (error bars) of the species that contributed the most to community dissimilarity in the four models: spring, summer, autumn and all (the three seasons combined).

Some environmental variables were log-transformed to achieve normality, in order to reduce skewed distribution of the response variable and hence reduce variance heterogeneity.

**Table 2.**
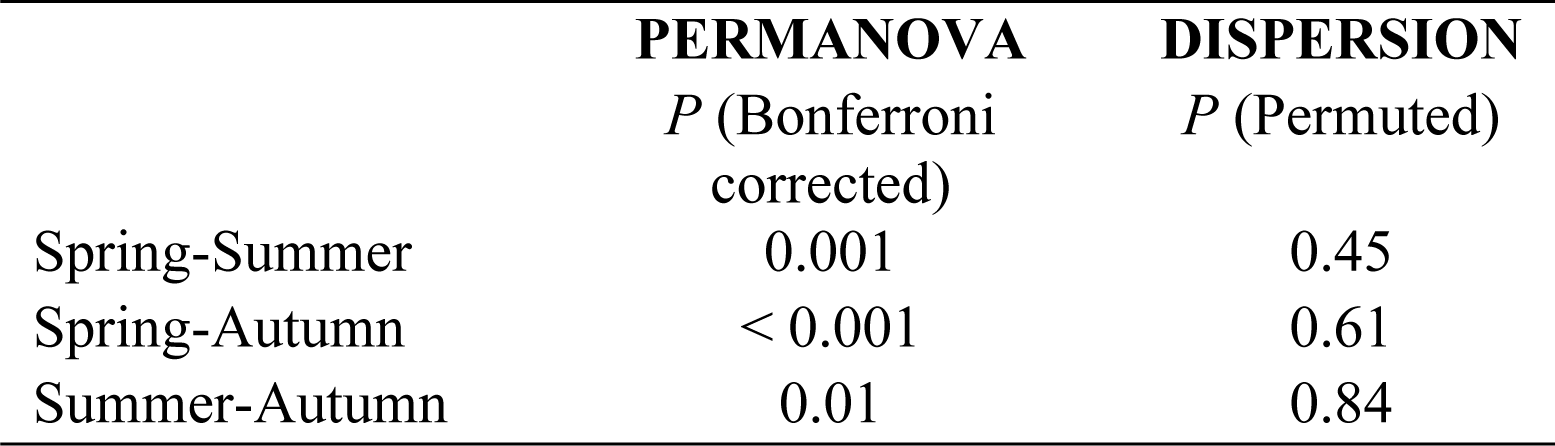
*P* values of PERMANOVA and test of multivariate dispersions comparing the three different datasets (spring, summer and autumn). Significance level of α=0.05.

**Table 3.** Species that contribute the most to community dissimilarity (based on Bray-Curtis measure) between the three sampling seasons (spring, summer and autumn). Average dissimilarity (Av. dissim), contribution percentage (contrib %) and cumulative percentage are shown.

| Taxon | Av. dissim | Contrib. % | Cumulative % |
| --- | --- | --- | --- |
| *Cocconeis placentula* | 10.21 | 12.28 | 12.28 |
| *Rhoicosphenia abbreviata* | 7.55 | 9.07 | 21.36 |
| *Tabularia fasciculata* | 3.94 | 4.74 | 26.10 |
| *Gomphonema parvulum* | 3.86 | 4.64 | 30.74 |
| *Nitzschia frustulum* | 2.98 | 3.58 | 34.33 |
| *Navicula perminuta* | 2.86 | 3.44 | 37.76 |
| *Ctenophora pulchella* | 2.74 | 3.29 | 41.06 |
| *Navicula recens* | 1.90 | 2.29 | 43.35 |
| *Diatoma tenuis* | 1.87 | 2.24 | 45.59 |
| *Navicula gregaria* | 1.79 | 2.16 | 47.75 |
| *Gomphonema olivaceum* | 1.76 | 2.11 | 49.86 |
| *Nitzschia fonticola* | 1.53 | 1.84 | 51.70 |
| *Mastogloia pumila* | 1.46 | 1.76 | 53.47 |
| *Diatoma moniliformis* | 1.27 | 1.52 | 54.99 |
| *Navicula tripunctata* | 1.09 | 1.31 | 56.31 |

If instead of using averages of environmental variables of the summer months we have used whole year averages of environmental variables, none of them was sufficient in explaining diatom composition in any of the individual seasonal datasets. Therefore, environmental variables were calculated per season and the CCA were produced for the corresponding seasonal diatom dataset. Salinity thereby exhibits the strongest explanatory power (λ1/λ2>1) for diatom community composition in the training dataset: λ1/λ2_spring_=1.16; λ1/λ2_summer_=1.29; λ1/λ2_autumn_=1.25 and λ1/λ2_all_=1.22 (Fig 4). Only two environmental variables showed statistical significant differences between the seasons according to One way ANOVA analysis: total nitrogen (F: 3.37; *P*=0.038) and pH (F: 6.76; *P*=0.002).

**Fig 4.**
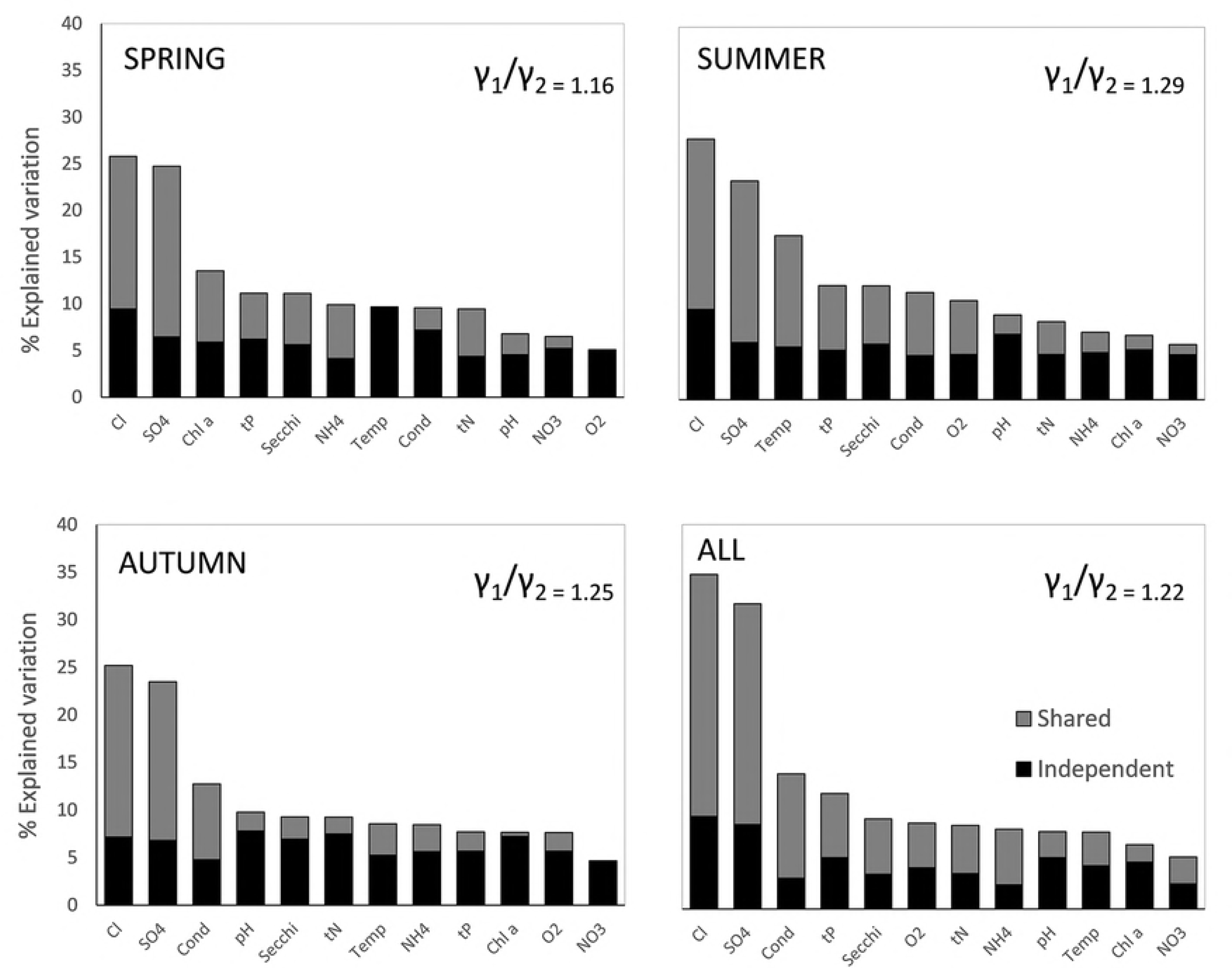
Variance partitioning of the training dataset (spring, summer, autumn and all combined). The independent and shared components of variance explained by the measured environmental variables and the ratio λ1/λ2 as a measure of the explanatory strength of salinity is also shown.

Hierarchical partitioning revealed that in all seasons sampled, salinity is the variable that explained most of the variation on diatom community composition. From the total explained variation, salinity explained a total variation of 25.8% in spring model (unique is 9.5%), 27.8% in summer model (unique is 9.7%), 25.2% in autumn model (unique is 7.2%) and 35% in ‘all-season’ model (unique is 9.7%) (Fig 4). The similar and high percentage of total explained variation by chloride and sulfates indicates the correlation between these two variables. Also relatively high and expected correlation is found with conductivity (Table 4). Because chloride correlate to sulfates and conductivity in a different degree according to the model, it was not possible to use any rule of thumb to discard collinear variables because otherwise, models cannot be compared. Therefore, we decided to keep all environmental variables measured in all models to see the overall pattern of variance partitioning and being able to compare the three models.

**Table 4.** Correlations between salinity and the rest of environmental variables in the different models and datasets (spring, summer, autumn, all seasons and validation). *P* values are shown for significant Pearson correlations. Temp= temperature; O_2_= Dissolved oxygen; tN= Total nitrogen; Tp= Total phosphorus; Secchi = Secchi depth; Cond= conductivity; Chl *a*= chlorophyll *a*.

|  | TEMP | O<sub>2</sub> | NO<sub>3</sub> | NH<sub>4</sub> | tN | tP | pH | SECCHI | COND | SO<sub>4</sub> | Chl<br>*a* |
| --- | --- | --- | --- | --- | --- | --- | --- | --- | --- | --- | --- |
| SPRING<br>n=32 | 0.15 | 0.06 | -<br>0.003 | 0.20 | -0.05 | 0.03 | -0.20 | -0.17 | **0.59**<br>**p<0.001** | **0.64**<br>**p<0.001** | -<br>0.14 |
| SUMMER<br>n=32 | **-0.54**<br>**p=0.001** | -0.33 | 0.02 | 0.12 | 0.24 | **0.36**<br>**p=0.04** | -0.16 | **-0.38**<br>**p=0.03** | **0.58**<br>**p<0.001** | **0.67**<br>**p<0.001** | 0.22 |
| AUTUMN<br>n=32 | **-0.4**<br>**p=0.02** | -0.34 | -0.01 | 0.21 | 0.02 | 0.08 | -0.12 | **-0.32**<br>**p=0.04** | **0.64**<br>**p<0.001** | **0.81**<br>**p<0.001** | 0.15 |
| ALL<br>n=96 | -0.16 | -0.19 | -<br>0.003 | 0.16 | 0.07 | 0.15 | -0.09 | **-0.26**<br>**p=0.01** | **0.60**<br>**p<0.001** | **0.70**<br>**p<0.001** | 0.10 |
| VALIDATION<br>n=90 | 0.07 | **0.27**<br>**p=0.01** | 0.41 | 0.23 | **0.36**<br>**p<0.001** | -0.08 | -0.09 | -0.08 | **0.97**<br>**p<0.001** | **0.54**<br>**p<0.001** |  |

The WA-PLS technique revealed that the most parsimonious models for salinity inference would be the one-component model for spring and autumn and the two-component model for summer and all seasons combined. The relationship between observed and predicted values of salinity were very strong with no apparent outliers (spring: r^2^jacknife=0.7; RMSEP=0.25; summer: r^2^jacknife=0.87; RMSEP=0.18; autumn: r^2^jacknife=0.78; RMSEP=0.26 and all: r^2^jacknife=0.86; RMSEP=0.19 Table 5; Fig 5).

**Table 5.** Performance statistics for weighted averaging partial least squares (WA-PLS) based diatom salinity inference models for spring, summer, autumn and all season models

| Transfer function | WA-PLS components | $r^2_{\text{(apparent)}}$ | $r^2_{\text{(Jackknife)}}$ | RMSE | RMSEP | $p$ values |
| --- | --- | --- | --- | --- | --- | --- |
| SPRING | 1 | 0.84 | 0.7 | 0.18 | 0.25 | <0.001 |
| SUMMER | 2 | 0.96 | 0.87 | 0.09 | 0.18 | 0.002 |
| AUTUMN | 1 | 0.90 | 0.78 | 0.17 | 0.26 | <0.001 |
| ALL | 2 | 0.92 | 0.86 | 0.14 | 0.19 | 0.004 |

The validation of the model using an independent dataset is shown in Fig. 6. The models to some extent overestimate salinity values at the low end of the gradients and underestimate the values at the high end, which was evident in the residuals *vs.* observed salinity plot (S1 Fig). The edge effect is clearly improved by the ‘all-season’ model, providing the most accurate reconstruction out of the four models (Fig 6).

**Fig 5.**
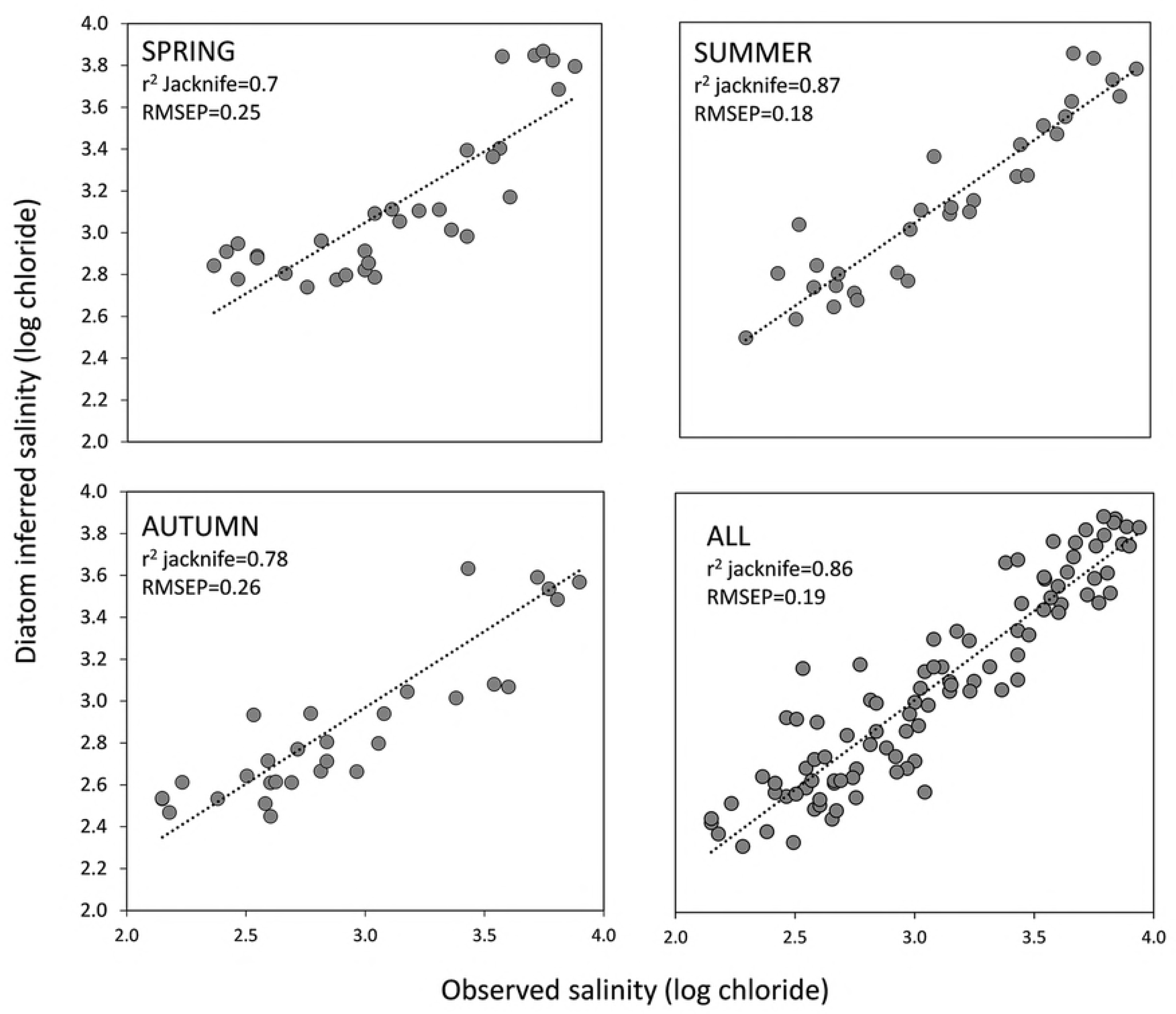
Diatom salinity inference models (WA-PLS) showing the relationship between observed and predicted salinity under leave-one-out cross validation for spring, summer, autumn and ‘all-seasons’ combined.

## Discussion

Our study shows that seasonality effects were responsible for important shifts in species dominance in our training set, as it is evident from the highly significant PERMANOVA test results. Changes in relative abundance of species during seasonal succession are usually advanced by the shift from early colonizers such as the adnate species *Navicula perminuta* [44] or poor competitors (*Ctenophora pulchella, Gomphonema olivaceum*) to species competing well under a resource limitation in late-successional communities (*Cocconeis placentula, Gomphonema parvulum*) [45-47]. Intra-species competition can be expected to play an important role in the nutrient-limited situation commonly occurring during late summer in freshwater and brackish environments [48]. Our data showed that total nitrogen significantly decreased in summer and autumn compared to spring. This could reflect changes in nutrient stoichiometry which in turn might be driving differences in seasonal species abundances. Besides competition under resource limitation, inter-specific interactions, such as grazing, are also important factors driving seasonal shifts in species abundances [13].

That seasonal changes of species abundances highlight the risk of developing transfer functions based on environmental variables measured in individual seasons, only. Despite the high number of studies highlighting the potential effect of confounding environmental variables and the application of temporal dependent calibrations [21, 49, 50] the effect of seasonality on the accuracy of diatom-based transfer functions has so far received only little attention. Calibration datasets based on samples taken during one season or calibrated against single measurement are very common [6, 51, 52] while in other cases sampling time is not specified [3, 42, 53]. In palaeocological studies, sediment core samples contains diatom assemblages that have already been subject to temporal and spatial integration. However, the development of the transfer function always required a modern data set. It is important to note that optima and tolerances of diatom species are different if the samples for developing the models are taken in spring, summer or autumn, even when salinity did not show statistically significant changes between the seasons (Fig.3). Especially important in this study was the contribution of *Cocconeis placentula* (with optima ranging from 1300 – 2800 mg/l chloride based on autumn and spring models respectively) or *Rhocoisphenia abbreviata* (with optima ranging from 900-1600 mg/l chloride based on autumn and spring models respectively) to seasonal community dissimilarity.

Potential inaccuracies and low predictive skills of models based on single season data from sites with highly variable hydrology and water chemistry were proposed earlier already [50]. For example when diatom calibration data sets use environmental measurements of annual means that do not necessarily reflect the specific concentrations present when diatoms are growing [54]. This mismatch may reduce highly significant variables to non-significant ones. As it is shown in our data, salinity became non-significant when annual averages were used.

The effect of seasonality investigated here have emphasized a crucial aspect negatively affecting the accuracy of transfer functions. That is, that almost in every environmental variable measured, an important fraction of explained variation is shared with other environmental variables [21]. In our study, even when the total percentage of explained variation of salinity is the highest in comparison to other variables, still around half of the total explained variation of salinity is shared with other environmental variables (sulphates, conductivity, secchi depth, total phosphorus and temperature). As it is expected, high correlation between salt ions (chloride and sulphates) and between ions and conductivity were found in the all training sets (spring, summer and autumn). Especially prone to be affected by secondary variables is the training data collected in summer and autumn where chloride concentration is significantly related to nutrient variables (total phosphorus), temperature and secchi depth (Table 4). In summer, phosphorus released from the sediment increases due to sulfate induced phosphorus mobilization [55], and this is reflected in the positive correlation between ions and total phosphorus. As a consequence of increased water nutrients, visibility is also affected. Both, the spring dataset and the combination of the three seasons, in the other hand, are less prone to the effect of secondary variables (e.g. nutrients) and showed lower correlation with potentially nuisance variables. The “all-season” model moreover showed the highest percentage of explained variation by salinity (Fig. 4). In addition the “all-season” model also improves the non-linear distortions at the end of the gradients (edge effects) that are an inherent problem of all unimodal-based calibration methods using weighted averaged estimations [36, 40, 49, 56] (Fig. 5). Although the weighted inverse deshrinking regression incorporated in WA-PLS reduces the edge effect, it inherently has its own problems by “pulling” the predicted values towards the mean of the calibration set resulting in an inevitable bias with some over-estimation at low values and some under-estimation at high values, as it is evident from S1 Fig [40, 49, 57]. In the “all-season” model this bias is reduced because the salinity signal is stronger (the total explained variation by salinity it is much higher than the variation explained by the other variables) and also because the number of samples has increased from 32 to 96 (which improved species optima estimation). The dataset size of the “all-season” model falls into the range of ≈ 100 suggested by Wilson [52] for the greatest improvement of the RMSEP. As a way of verification if the good performance of the “all-season” model is mainly due to the higher number of samples in comparison to the other models, we also built the all-season model by randomly selected one sample out of the three seasons in all sampling locations. The result of the model performance is the same: r^2^jacknife=0.86; RMSEP=0.19.

When we tested the performance of the models using the completely independent validation dataset, we also found that the “all-season” model performed the best. Although the four models performed relatively well in the model cross-validation, the independent validation demonstrate the importance of a comprehensive dataset that takes on board the differences in species abundances due to seasonality, as in our “all-season” model. The independent validation confirms the improvement of the non-linear distortions. The underestimation of the high salinity values is the lowest in the “all-season” model. As a result, this model provides the most realistic reconstruction even though some edge effects remain (Fig 6).

**Fig 6.**
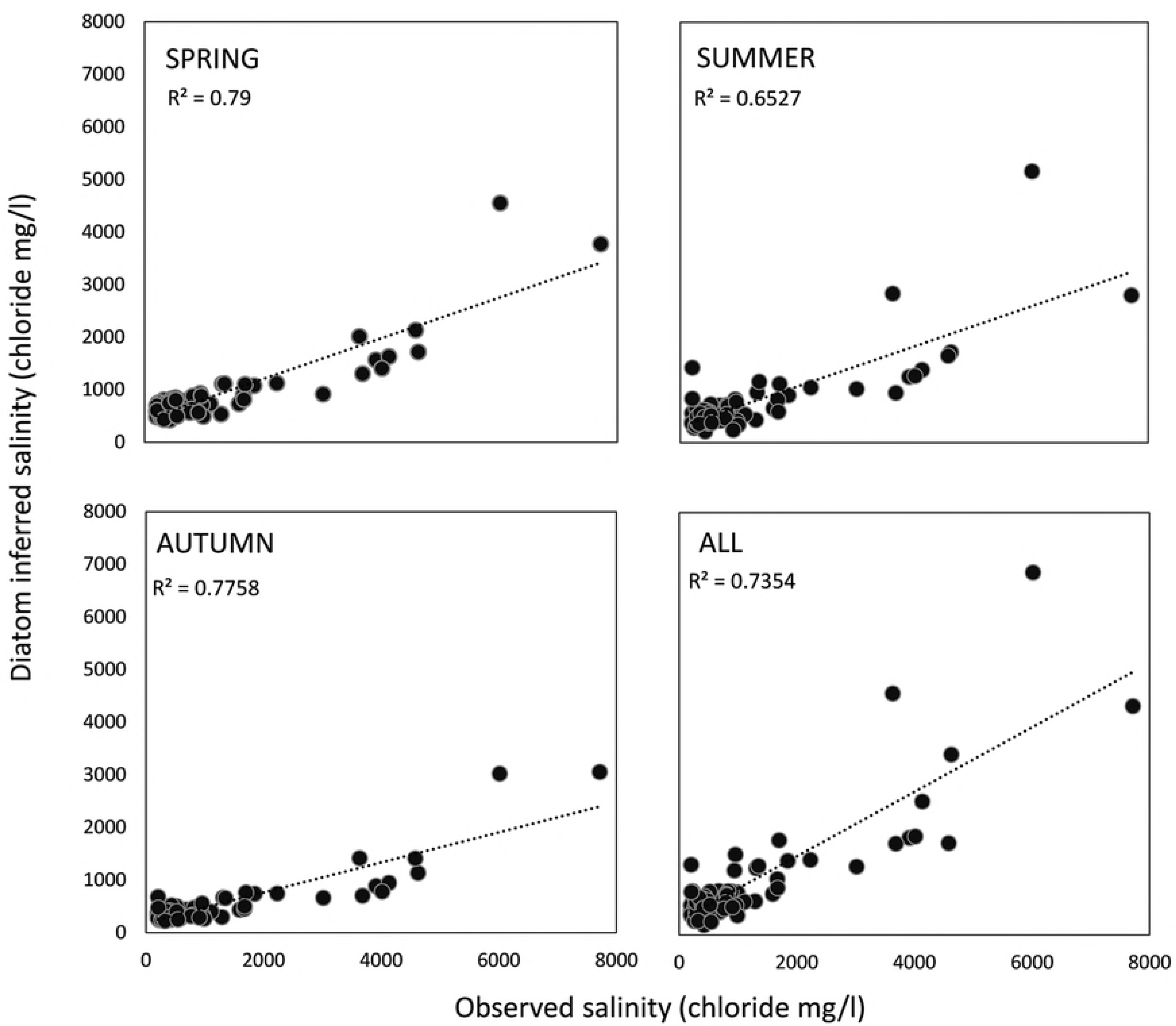
Salinity reconstructions using the validation dataset for spring, summer, autumn and ‘all-seasons’ WA-PLS model.

The reasons of this offset are several: First, the effect of secondary variables in the training sets. The significant correlation between salinity and potential nuisance variables in summer and autumn (total phosphorus, secchi depth and temperature) lies between 0.32-0.54 R^2^ values (Table 4). According to Juggins (2013), for training sets that exhibit a correlation 0.2< R <0.5 to a nuisance variable, a change in the latter can produce spurious fluctuations in the reconstructed variable typically greater than 10 units. In addition, even a low correlation to a nuisance variable (R <0.2) affects the reconstruction in at least 10 units. This corresponds to 0.1 of the gradient length studied in Juggins [21] which in this study will be equivalent to ≈880 mg/l of chloride change. In the independent validation we have found a bias of less than 500 mg/l chloride in around 68%-77% of the observations in the four models and a bias between 2000-4000 mg/l chloride in around 12% (autumn model) - 4.5% (“all-season” model) of the observations. Second, the ratio λ1/λ2 in the validation dataset is lower than in the models (λ1/λ2_validation_ =0.88; S2 Fig). The significant correlation between salinity and nuisance variables in the validation dataset (total nitrogen and dissolved oxygen- Table 4) could be the reason of this lower ratio and effectively alter the driving effect of chloride in the distribution of diatom communities. When transfer functions are used to performed palaeoenvironmental reconstructions, the strength of the relationship between the species and the modelled variable is not known. It is therefore highly important that the transfer function is based on a comprehensive dataset covering seasonal variability and with a sample size of ≈100 to reduce the model bias as much as possible, since it is not possible to reduce the noise of the palaeo data. Third, there are always some species mismatches between the transfer function and the reconstruction data which lessen the accuracy of palaeoenvironmental reconstructions. In our study, indicators of high salinity in the models (*Mastogloia pumila, Rhopalodia brebissonii, Navicula arenaria*) were not present or very scarce in the validation dataset. This could be due to problems in the taxonomic identification (e.g. species of the genus *Navicula*), differences in the number of individuals identified between the model and the reconstruction datasets, or due to ecological factors not taken into account (dispersal, grazing, etc). In addition, brackish species usually have very wide tolerances due to the fact that these species are well adapted to the fluctuating environmental conditions in shallow brackish habitats [58]. An example are species of the genus *Diatoma* [59]. Other species such, *Rhoicosphenia abbreviata* and *Cocconeis placentula* were present in the whole salinity gradient. These are pollution tolerant species [60] that do not respond to the salinity changes considered in this study and thus, may also introduce noise in the reconstruction values. This could be the reason why the greatest bias in model performance occur in the high end of the gradient. High salinity observations (>2000 mg/l chloride) represents only 12% of all the samples in the validation dataset and euhalobe or polyhalobe diatoms may be underrepresented. With the general aim to collect representative community data, not the taxonomic identification but also the quality of the indicator may have a considerable impact in (palaeo) environmental reconstructions. Species that have very wide tolerances to the environmental variable of interest (for chloride they are called euryhaline forms) may therefore be eliminated for transfer function models. This is a very interesting field of research and deserves further investigation, so that appropriate recommendations can be made towards further standardization.

## Conclusions

Assuming that diatom inference models are invariant in space and time has important consequences for the reliability of environmental reconstructions. The present study has shown that seasonality controls the predictive skills of diatom transfer functions by driving changes in the abundance of the dominant species. While transfer function performance of the single season models (spring, summer or autumn) is reasonably good in the model cross-validation, the validation of the models with an independent dataset demonstrate that single season sampling leads to a strong underestimation of salinity due to the confounding effect of other environmental variables such as temperature, secchi depth, nutrients and dissolved oxygen. The strength of salinity as the main environmental variable explaining the distribution of diatom communities is enhanced in the “all-season” model. The “all-season” model combines all data and provides the highest percentage of explained variation by salinity, increases the accuracy of species optima and tolerance estimations, and reduces the non-lineal distortions at the end of the gradients.

One of the basic requirements for quantitative palaeoenvironmental reconstruction is the availability of a large, high-quality data set of modern species assemblages and contemporary environmental data. Given the strong effect of seasonality on the composition of diatom assemblages, the development of transfer functions capturing intra-annual variability of species abundances and independent model validation is strongly recommended.

## Acknowledgements

This study was financed by Utrecht University. Special thanks to Gert van Ee (*Hoogheemeraadschap Hollands Noorderkwartier*) and Herman van Dam (*Water en Natuur*) for providing the data set used to build the models and *Hoogheemraadschap Hollands Noorderkwartier* for providing the ecological monitoring data (validation dataset).

## Supporting information

**S1 Fig. Residuals vs observed salinity with corresponding R^2^ values in the four models: spring, summer, autumn and all season models.** The overestimation of salinity at low end and an underestimation of salinity at the high end of the gradient is especially evident in the spring and autumn models. When all seasons are combined, this edge effects are less prominent and a more accurate performance results.

**S2 Fig. Comparison of the percentage of unique and shared explained variation by salinity in the different datasets (spring, summer, autumn, all seasons and validation datasets).** The ratio of λ1/λ2 is also shown for each case.

